# Reliable detection of pyrazinamide antitubercular activity *in vitro*

**DOI:** 10.1101/2022.05.22.492909

**Authors:** Alexandre Gouzy, Claire Healy, Dirk Schnappinger, Sabine Ehrt

**Affiliations:** Department of Microbiology and Immunology, Weill Cornell Medical College, New York, NY 10065, USA

**Keywords:** Pyrazinamide, Mycobacterium tuberculosis, Lipids, acidic pH, drug resistance

## Abstract

Pyrazinamide (PZA) is a pivotal antibiotic for the chemotherapy of tuberculosis (TB), a disease caused by the bacterium *Mycobacterium tuberculosis* (Mtb). PZA is notorious for its poor *in vitro* activity which complicates phenotypic PZA-susceptibility testing (PPST) and likely causes inappropriate treatment of TB patients. Here, we show that PZA activity can be reliably detected using an acidic and lipid-rich culture medium mimicking conditions Mtb encounters during an infection. Our growth model could facilitate PPST to improve the treatment of TB patients and ameliorate the global surveillance of PZA resistance.

## Introduction

Tuberculosis (TB) is caused by *Mycobacterium tuberculosis* (Mtb) and is the deadliest human disease due to a single bacterium (WHO, Global TB report, 2020). The rise of drug resistance is a major public health concern for the treatment of infectious diseases such as TB [1]. Pyrazinamide (PZA) is a pivotal first-line anti-TB drug that allowed the shortening of TB treatment from 9 to 6 months [2]. PZA resistance is estimated to affect 3 to 42.1% of TB cases and is mainly due to mutations in the promoter or the open reading frame of the gene *pncA* (80-90%) coding for the enzyme PncA [3], [4]. PncA converts PZA into its active form, pyrazinoic acid (POA). The mechanism(s) of action of PZA and POA are still not fully elucidated [5]. DNA sequencing of the *pncA* region is currently the most reliable method for the detection of PZA resistance [6] but the technology required for such tests is not universally available in area of the world where TB is prevalent. Moreover, the absence of a “hot spot region” comprising most mutations in the *pncA* region complicates the development of a diagnostic assay [4]. Therefore, the World Health Organization considers phenotypic PZA-susceptibility testing (PPST) the gold standard for the detection of PZA-resistant TB. Despite its excellent efficacy *in vivo*, PZA is notorious for its poor activity against Mtb in standard media making PPST technically challenging. PZA activity increases in acidic media and commercial PZA testing kits use media buffered to pH5.9 to detect PZA resistance. The reliability of PPST in the clinic is reduced by the high frequency of false positive results caused by large inoculum sizes increasing the pH of the media to a pH at which PZA is not active [7]. Moreover, growth of Mtb is inhibited at pH<5.8 in standard media containing glycerol (Gly) and glucose as main carbon sources, which prevents the use of media at pH lower than 5.9 [8]. Consequently, PPST is not routinely conducted in the clinic which prevents optimal treatment of patients infected with PZA-resistant Mtb [9]. We have recently shown that Mtb can readily grow in acidic media (pH<5.8) if host-relevant lipids such as oleic acid (OA) are provided as the primary carbon source [8]. Here, we demonstrate that an acidic and lipid-rich culture medium allows to accurately measure the minimal inhibitory concentration (MIC) of PZA to detect PZA resistance.

## Results and discussion

To quantify PZA activity *in vitro*, wild-type Mtb was grown in 7H9 media containing Gly or OA as a main carbon source at either neutral (pH7) or acidic pH (pH5) (Fig 1 *A*). 100 μg/mL PZA did not affect growth of Mtb at pH7 and as previously described, Mtb did not grow with Gly at pH5 independently of the presence of PZA [8]. In contrast, 100 μg/mL PZA completely inhibited the growth of Mtb with OA at pH5. Moreover, PZA drastically reduced Mtb CFUs at pH5 in the presence of OA (>6 log10) but only had a minor impact with Gly (≈1 log10) at pH5. Furthermore, by growing Mtb with OA at various pH, we demonstrated that decreasing the pH increases Mtb growth inhibition and killing by PZA (Fig. 1 *B* and *C*). In these experiments, growth with OA depended on OA replenishment, which was needed to avoid the toxic effect OA exerts on Mtb at high concentrations. While this OA replenishment method stimulates growth of Mtb, it limits throughput. Early studies by René Dubos have shown that bovine serum albumin (BSA) binds to OA and protects Mtb from OA toxicity [10]. To incorporate more OA in our media without causing Mtb growth inhibition, we increased the amount of BSA usually present in the media (50g/L instead of 5g/L). In this BSA-enriched medium, Mtb grew with up to 7 mM OA at pH7 and 4 mM at pH5 (Fig. 1 *D*). The increased OA toxicity observed at acidic pH is likely due to the reduced binding of OA to BSA leading to higher amounts of free toxic OA [11]. Using BSA-enriched media containing 3 mM OA, we determined the MIC of PZA at various pH (Fig. 1 *E*). Bacterial growth was inhibited by less than 25% at ≥pH 6.5 and less than 65% at pH6.25 with the highest concentration of PZA tested. In comparison, PZA fully inhibited Mtb growth at pH6 and lower (MIC90pH6= 534 μg/mL; MIC90pH5.75= 594 μg/mL; MIC90pH5.5= 392 μg/mL; MIC90pH5.25= 137 μg/mL; MIC90pH5= 79 μg/mL). The MIC90 of PZA measured in our growth model are similar to those reported in previous studies [12], [13]. Moreover, we confirmed that PZA activity is increased at acidic pH as described in early *in vitro* studies and more recently in the macrophage model of infection [12], [14]. As a proof of concept for PPST, we tested the resistance to PZA of an Mtb strain lacking *pncA*. Using OA replenishment at pH5, we first showed that growth and survival of the *ΔpncA* strain were not much affected by 500 μg/ml PZA (Fig. 2 *A* and *B*). Using a BSA-enriched medium at pH5.5 containing 3 mM OA, we reliably detected high-level resistance of the *ΔpncA* mutant to PZA in comparison to its WT and the complemented mutant (Fig. 2 *C*). In accordance with the role of PncA in converting PZA into POA, the *ΔpncA* mutant strain was as sensitive as WT to POA (Fig. 2 *C*). The detection of PZA resistance was shown to improve patient outcome [9]. Thus, the culture medium developed here could help implement PPST routinely in the clinic and ameliorate patient care. As PZA is currently considered to be part of a novel anti-TB regimen, an improvement of PPST might be beneficial for TB treatment now and in the future [15].

**Fig. 1.**
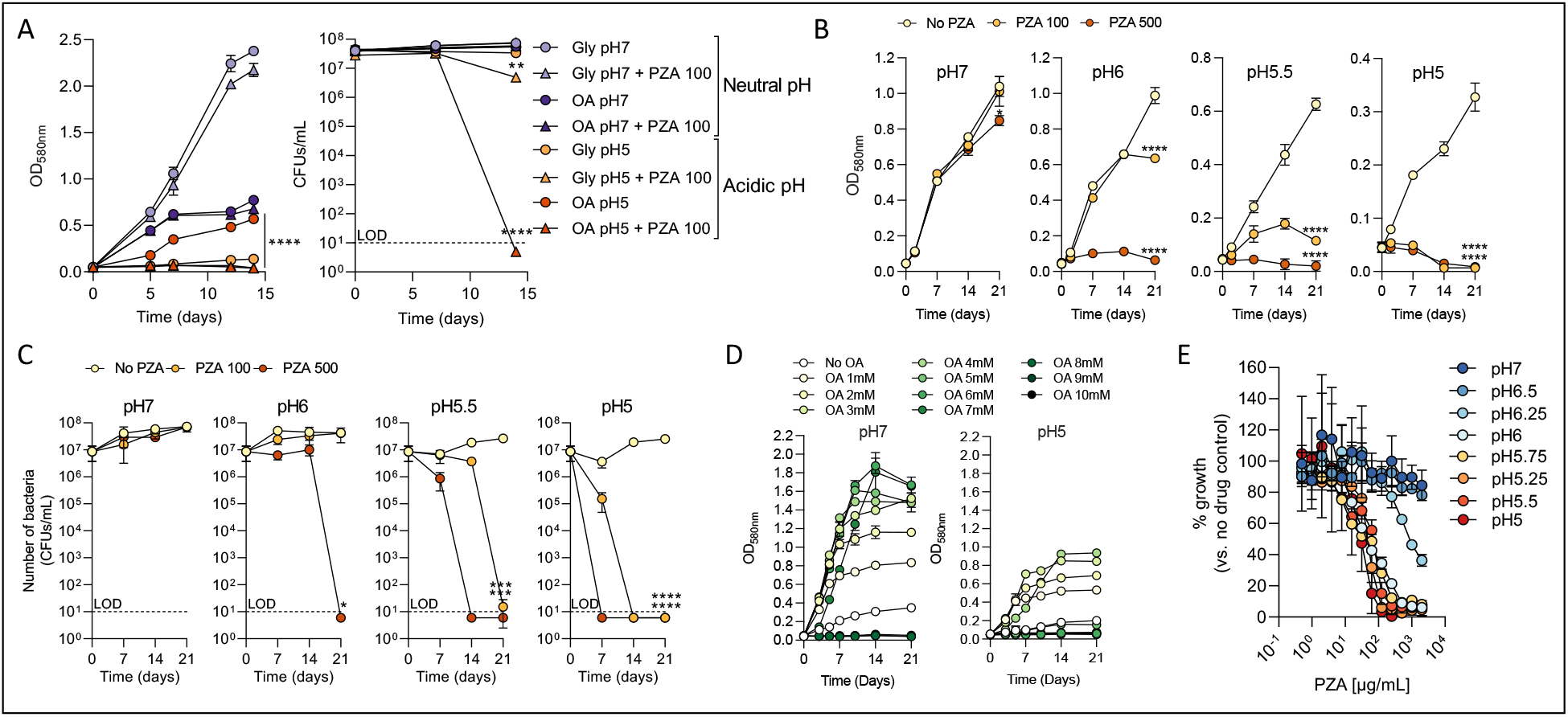
Antitubercular activity of PZA in lipid-rich and acidic conditions. *(A)* Growth (left panel) and survival (right panel) of WT Mtb in 7H9 with Gly (20mM) or OA at either pH7 or pH5 and in the presence or not of PZA (100 μg/mL). Growth (*B*) and survival (*C*) of WT Mtb at various pH with OA and in the presence or not of PZA at 100 or 500 μg/mL. (*D*) Growth of Mtb at pH7 and pH5 in 7H9 containing BSA at 50g/L and various amount of OA. *(E)* Growth of Mtb at various pH and against various concentrations of PZA in 7H9 containing 3mM OA and 50g/L of BSA. Data were recorded after 10 days at 37°C. In A-C, 200μM OA was replenished ev. 2-3 days [8]. In A-C, statistical significance was determined for the final time points by comparing PZA-treated conditions with control samples not containing PZA using a one-way ANOVA and Tukey’s multiple comparison test. Data in A-D are means ± SD of three independent experiments and data in E are means ± SD of an experiment performed in triplicate and representative of three independent experiments. LOD: Limit of detection. CFU: colony forming unit. *\*P*< 0.05; \*\**P*< 0.005; \*\*\**P*< 0.0005; \*\*\*\**P*< 0.0001.

**Fig. 2.**
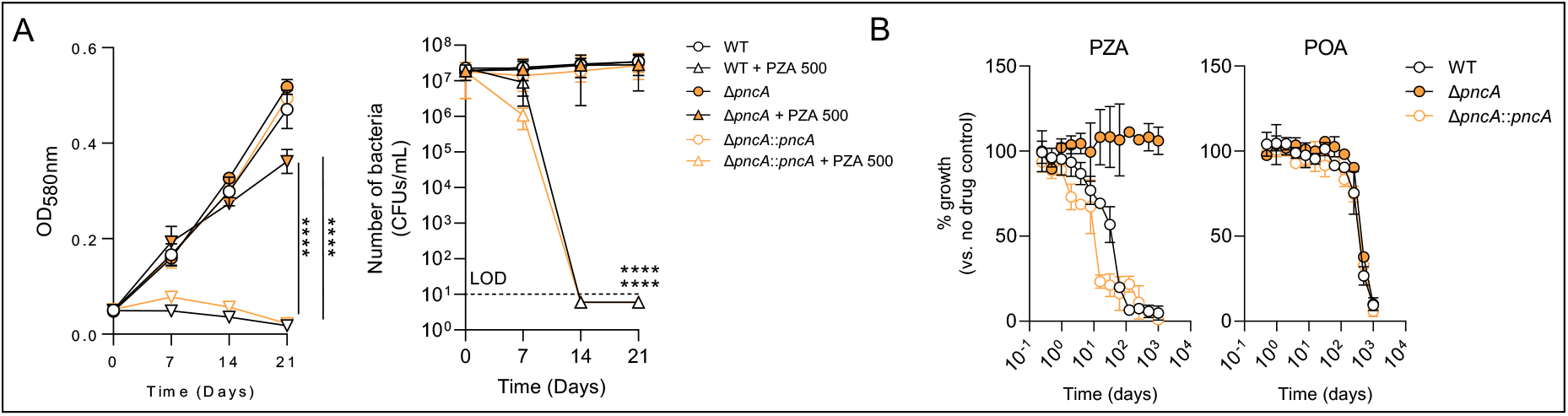
Reliable detection of PZA resistance *in vitro. (A)* Growth (left panel) and survival (right panel) of wild-type (WT), *pncA* mutant (Δ*pncA*) and complemented strain (Δ*pncA*::*pncA*) at pH5 with OA in the presence or not of PZA at 500 μg/mL. (*B*) Growth of WT, *ΔpncA* and Δ*pncA*::*pncA* strains at pH5.5 against various concentrations of PZA or POA in 7H9 containing 3mM OA and 50g/L of BSA. Data were recorded after 10 days at 37°C. In A, 200μM OA was replenished ev. 2-3 days [8]. In A, statistical significance was determined for the final time points and among PZA-treated conditions using a one-way ANOVA and Tukey’s multiple comparison test. Data in A are means ± SD of three independent experiments and data in B are means ± SD of an experiment performed in triplicate and representative of three independent experiments performed in triplicates. LOD: Limit of detection. CFU: colony forming unit. \*\*\*\**p*< 0.0001.

## Materials and Methods

The wild-type Mtb strain used was H37Rv. In Fig. 2 parental H37Rv wild-type and *ΔpncA* mutant strains were used along with the *ΔpncA* mutant complemented by chromosomal (attL5) expression of *pncA* under the control of its native promoter. Prior to growth experiments, bacteria were grown in liquid culture in complete Middlebrook 7H9 medium supplemented with 0.2% glycerol, 0.05% tyloxapol, and ADN (5 g/L BSA, 0.2% dextrose and 0.085% NaCl). Before growth experiments, bacteria were washed twice in phosphate buffer saline and resuspended in 7H9 base media + 0.05% tyloxapol + 0.085% NaCl containing either 5g/L or 50g/L of fatty acid free BSA (fraction V Roche) with no glycerol nor dextrose added. Media were supplemented with glycerol (20mM) or OA at various concentration from stocks at 1% (40 mM) or 3% (120mM) OA. pH adjustment of media was carried out by adding HCl or NaOH when appropriate. MES (2-(N-Morpholino) ethanesulfonic acid) hydrate was added to acidified media (pH≤6.5) at 100 mM. In Fig. *1E*, MIC90 were calculated using the Gompertz equation. For survival experiments, bacteria were plated onto 7H10 plates containing 4g/L of charcoal.

## Acknowledgments

We thank Helena I. M. Boshoff for providing the *Mycobacterium tuberculosis* H37Rv *ΔpncA* mutant strain and its wild-type counterpart. This work was supported by the NIH Grant No. P01AI143575.

